# A neural mechanism for online discovery of latent contexts

**DOI:** 10.64898/2026.03.31.715618

**Authors:** Ali Hummos, Mien Brabeeba Wang, Qihong Lu, Kenneth A. Norman, Mehrdad Jazayeri

## Abstract

Experience unfolds as a stream shaped by hidden causes that change over time. Adaptive behavior requires inferring the underlying states and adjusting when they change. Yet, how neural circuits discover and track latent states remains unclear. Here we introduce NeuraGEM, a neural architecture that combines fast transient activity with slow synaptic plasticity to implement an online analogue of Expectation–Maximization. By separating timescales, NeuraGEM clusters sequential experiences, detects context changes, and stabilizes task-specific computations. The model generalizes beyond conventional recurrent networks and reproduces key features of human contextual learning, including curriculum-dependent effects. It also gives rise to population dynamics resembling those observed in brain circuits, including line-attractor structure and transient error responses at change points. Together, these findings provide a mechanistic account of how neural circuits organize experience into latent states that support rapid inference and adaptive behavior.

---

Our experiences unfold as a continuous stream, yet subtle external changes may signal important changes demanding a shift in behavior. For example, in a conversation, the words, facial expressions, and gestures may remain largely unchanged from one moment to the next. Yet, a slight change of intonation or a long pause can mark a shift in the partner’s internal state—from curious to impatient, from engaged to withdrawn (fig. 1). In essence, we can infer latent contexts remarkably efficiently from noisy, ambiguous observations. However, the computational mechanisms that underlie this capacity are not known.

**Fig. 1.**
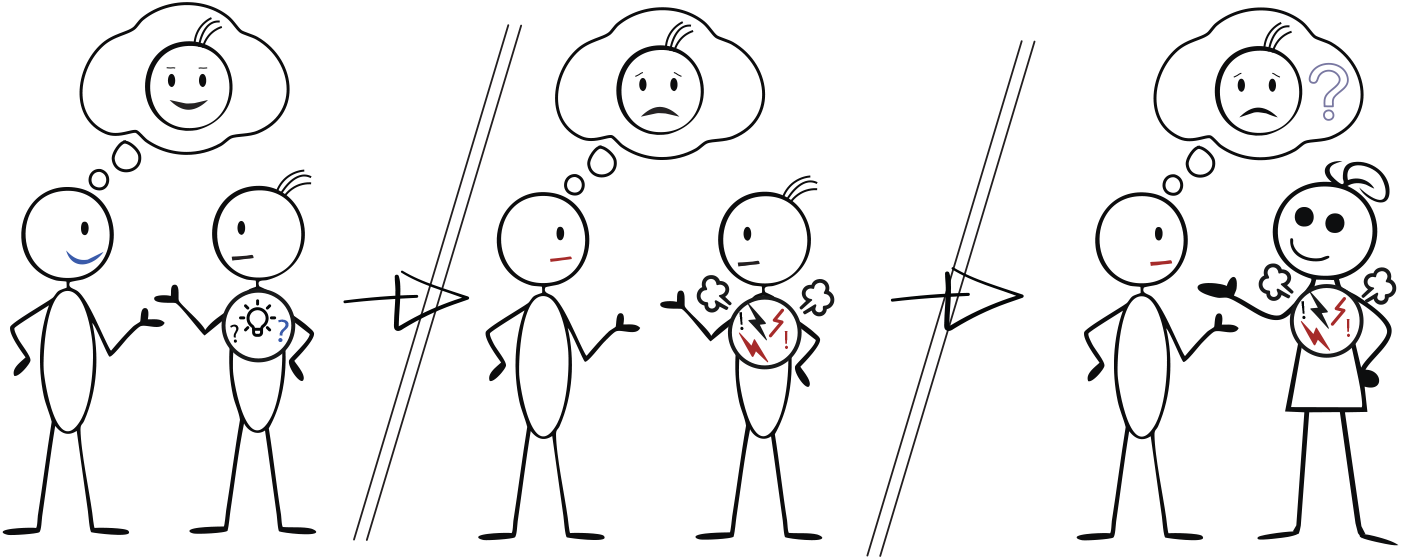
A schematic of learning a categorization of emotional states in others. Others’ emotional states are a latent variable that we have limited access to. Discovering the existence of such latent states allows us to adapt our behavior and facilitates generalization to new situations.

Numerous studies have examined how the brain organizes behavior according to inferred latent contexts [1–10] and generalizes newly learned relationships to novel conditions [3, 7]. Cognitive models capture these abilities through probabilistic inference over latent variables, highlighting context learning as a fundamental feature of adaptive behavior [1, 7, 11]. Yet, these models largely describe the computation in terms of how probabilities should be updated without specifying how neural dynamics and plasticity enable making inferences in real time.

Neural recordings have identified correlates of hidden contexts in the frontal and prefrontal cortices. A common observation is that neurons in the prefrontal cortex encode the current context, while those in the anterior cingulate cortex track prediction errors and accumulate evidence for context switches [12–18]. However, since most studies have focused on well-trained animals, we do not know how neural circuits rapidly detect and flexibly adapt to new latent contexts.

Computational approaches have offered partial insights. Recurrent neural networks (RNNs) can, in principle, represent hidden states, but they typically rely on slow, weight-based learning and suffer interference when latent states change [19–22]. RNNs trained with long input horizons that span many context switches can approximate Bayesian inference over contexts [23]; however, they require massive datasets and mechanisms for credit assignment over extensive time spans, limiting biological relevance. At the other extreme, switching linear dynamical systems [24] or hierarchical hidden Markov models [25, 26] often make a priori assumptions about latent states, leaving unanswered how latent context representations arise naturally in neural systems.

Here, we introduce NeuraGEM, a simple neural architecture that learns latent contexts online through local, error-driven updates. NeuraGEM comprises two interacting neural modules operating at two distinct timescales [22, 27]. One module functions as a leaky integrator and enables rapid adjustments to sudden errors, while the other slowly learns the task to lower overall errors. The two modules complement each other in enabling rapid and flexible adaptation to latent context switches while minimizing interference during learning. We demonstrate that the interactions between the two modules establish a neural analogue of the Expectation–Maximization algorithm. This mechanism enables the system to dynamically cluster experiences, detect boundaries between regimes, and form compact context representations that generalize beyond the training data. Notably, NeuraGEM reproduces key behavioral features of human contextual learning, matches neural signatures of context representation and switching, and offers a mechanistic framework for how the brain discovers latent structure from the statistical regularities of experience.

## 1 Results

We begin with a simple one-dimensional prediction task featuring latent contexts. On each trial, a sample is drawn from a Gaussian distribution whose mean alternates covertly between 0.2 and 0.8 (s.d.= 0.3). The mean alternations occur unpredictably in blocks of 15–50 trials (truncated geometric distribution, mean 25; fig. 2A). The task is to predict the next sample as accurately as possible based on past observations.

**Fig. 2.**
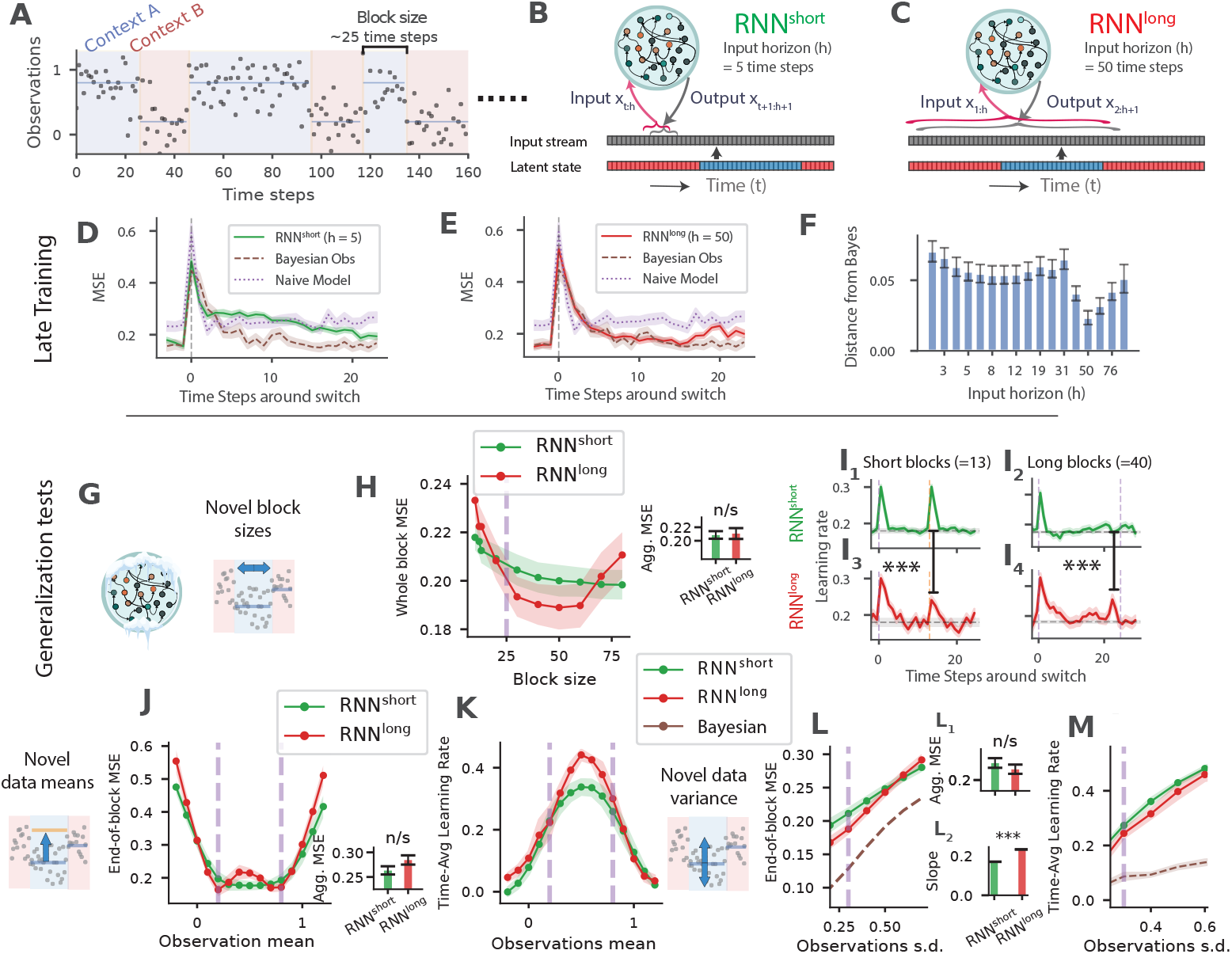
Conventional RNNs showed limited generalization. **A**) Example task observations. Observations were drawn from a Gaussian distribution with mean switching covertly between 0.2 and 0.8 (s.d. 0.3). Block lengths were sampled from a truncated geometric distribution (range 15–50 time steps; mean 25). Background shading indicates latent context. **B, C**) Input horizon *h* defines the number of recent samples available to produce output at each time step. Models updated weights every time step, and at the next time step processed the last *h* inputs again. **D, E**) Switch-aligned mean squared prediction error (MSE) for short-horizon RNNs (RNN^short^) (**D**) and long-horizon RNNs (RNN^long^) (**E**). Traces show error averaged across model seeds and switches late in training (blocks 30–40). The Bayesian observer uses the generative model underlying the task, whereas the naive model outputs the mean of the two most recent predictions (see Methods section 3.3.2). Shaded regions denote s.e.m. across 20 model seeds. **F**) Distance between model predictions and those of an ideal Bayesian observer, computed as the L1 norm (mean absolute deviation) across time, for RNNs with input horizons ranging from 1 to 100. Bars show mean deviation *±* SEM, n=20. **G**) Schematic indicating frozen weights during testing. **H, I**) Generalization to unseen block lengths. **H**: Mean squared error is averaged across time steps within each block. Average block size during training is indicated by the dashed purple line. Inset: Aggregate time-averaged MSE across block sizes (paired t-test, n = 20 independent model runs): RNN^short^ (mean *±* s.e.m. 0.209 *±* 0.0056) vs RNN^long^ (0.211*±*0.008), *t*(19) = −0.24, *p* = 0.82. **I**: Behavioral learning rate—defined as the fraction of the previous prediction error corrected on the subsequent prediction—shown for short and long blocks for RNN^short^ (upper panels, I_1_, I_2_) and RNN^long^ (lower panels, I_3_, I_4_). For short blocks, the difference in mean behavioral learning rate at the post-switch time step (*t* = 14) was Δ = 0.072 with *SE*_Δ_ = 0.020, *z* = 3.61 (normal approximation based on SEMs), two-sided *p <* 0.001. For long blocks, the difference at *t* = 23 was Δ = −0.114 with *SE*_Δ_ = 0.032, *z* = −3.58, two-sided *p <* 0.001. **J, K**) Generalization to unseen latent means. **J**: MSE when testing on Gaussian means between −0.2 and 1.2 (trained means 0.2 and 0.8 indicated by dashed lines). Inset: Aggregate time-averaged MSE across mean values (paired t-test, n = 20): RNN^short^ (0.263 ± 0.0083) vs RNN^long^ (0.285 ± 0.0094), *t*(19) = −1.63, *p* = 0.12. **K**: corresponding behavioral learning rate. **L, M**) Effects of increased observation noise. **L**: Prediction error as noise standard deviation is increased beyond the training level (*σ >* 0.3), compared to a Bayesian model extended to infer sample variance from the data. Inset L1: Aggregate time-averaged MSE across noise levels (paired t-test, n = 20): RNN^short^ (0.23 ± 0.0085) vs RNN^long^ (0.219 ± 0.0081), *t*(19) = 0.93, *p* = 0.36. Inset L2 shows the slope of error growth for each model. Slopes differed significantly between models (RNN^short^: 0.17 *±* 0.00058; RNN^long^: 0.235 *±* 0.00058; two-sided paired t-test, *p <* 10^−6^). **M**: Behavioral learning rate under the same noise manipulations.

### 1.1 Conventional recurrent neural networks

To illustrate the computational challenge of context inference, we first examined the performance of conventional recurrent neural networks (RNNs) trained on this task. A key parameter in RNN training is the input horizon *h*; i.e., the number of previous samples to which the model has access during training. This variable plays a crucial role in performance. When the input horizon is short, the training set does not furnish the opportunity to observe the block structure, and the model adapts to local observations mostly from one context. In contrast, when the input horizon is sufficiently long, RNNs can learn and adapt to the block structure more effectively.

We examined RNNs in these two regimes: a short horizon regime (*h <* 25), which, at best, exposes the network to one switch (fig. 2B), and a long horizon regime (*h >* 25) that could include multiple switches (fig. 2C). We refer to the former as RNN^short^, and the latter as RNN^long^.

RNN^short^ models were able to perform the task with moderate success but were slow to adapt after each switch. In these networks, each switch led to a sharp increase in error, followed by decay that was much slower than the ideal Bayesian observer (fig. 2D). Notably, this pattern persisted even late in training, indicating that these models did not retain information about both distributions and instead had to relearn the relevant statistics after each switch (i.e., catastrophic forgetting).

RNN^long^ models, in contrast, adapted more effectively. Following a switch, their errors declined rapidly, closely matching the performance of an ideal Bayesian observer (fig. 2E). This pattern indicates that these models retained information about both latent mean values and did not depend on updating RNN synaptic weights after each switch. This finding aligns with previous studies showing that longer input horizons facilitate meta-learning [23, 28].

We next assessed the models’ ability to generalize to novel task parameters. To this end, we froze the network weights and evaluated performance under new conditions, including unseen block sizes, novel sample means, and increased sample noise. For RNN^long^ models, we used an input horizon of *h* = 50, which produced the closest correspondence to the ideal observer (fig. 2F).

#### 1.1.1 Generalization to block length

We first quantified the performance of the models as we varied the block length. RNN^short^ models perform poorly after block switches (fig. 2D). Consequently, their overall performance was worse for shorter blocks with more frequent switches (fig. 2G,H). In contrast, RNN^long^ models exhibited a U-shaped performance profile: they performed best at intermediate block lengths—the range encountered during training—but poorly on both shorter and longer blocks (fig. 2G,H), indicating overfitting to the training distribution.

To probe these characteristics more closely, we computed an effective trial-by-trial *learning rate*, defined as the change in model output relative to the previous prediction error (See Methods section 3.3.1). Ideally, this learning rate should remain low within a block and increase only when there is evidence of a block switch.

RNN^short^ models showed a sharp increase in learning rate following block switches, and this transient response was largely insensitive to block length (fig. 2I, top panels). In contrast, RNN^long^ models displayed signs of overfitting to the block sizes experienced during training. When tested on shorter blocks, learning rate had a diminished peak after the unexpected switch point, while on longer blocks, learning rate rose near the *expected* rather than the *actual* switch point. In other words, RNNs^long^ under-reacted to early switches and over-reacted around the trained block duration, effectively hallucinating switches that did not occur (fig. 2I, lower panels).

#### 1.1.2 Generalization to new latent mean values

Next, we evaluated generalization to novel mean values for the input observations. Both RNN^short^ and RNN^long^ models performed well when latent mean values were close to those encountered during training (fig. 2J). However, performance declined markedly as the mean values departed from the training range. Specifically, both models showed progressively worse performance for mean values below 0.2 or above 0.8 (fig. 2J). Moreover, RNN^long^ models exhibited signs of overfitting to the training means, with performance also deteriorating for intermediate values within the 0.2–0.8 range (fig. 2J). This decline was mirrored in a complementary analysis of learning rate, which revealed increased RNN^long^ learning rate for ambiguous mean values between 0.2 and 0.8 (fig. 2K).

#### 1.1.3 Generalization in the presence of higher variability

Finally, we tested the models’ performance in the presence of larger sample variance. As a baseline, we used a Bayesian model that estimated observation noise from the data. Larger sample variance increased error in RNNs beyond the Bayesian model (fig. 2L), but RNN^long^ was more sensitive and had a steeper increase in error compared to RNN^short^ (fig. 2L, inset 2). The higher error in RNNs is due to the inflation of their learning rates, while the Bayesian model adjusted the sample noise estimate and maintained a lower learning rate (fig. 2M).

Together, these results demonstrate that conventional RNNs can successfully perform this simple prediction task and, when provided with long-horizon inputs, can approximate near-optimal inference. However, their performance deteriorates in generalization tests, indicating that they overfit to the latent statistics encountered during training and fail to generalize beyond them.

### 1.2 NeuraGEM

The behavior of conventional RNNs revealed a tension between two modes of learning (fig. 2). RNNs trained with short input horizons learn slowly because they must update synaptic weights—a process that unfolds in a large parameter space, which maintains adaptability but is vulnerable to interference and forgetting. Extending the input horizon allows RNNs to leverage meta-learning, effectively shifting part of the learning into the *activity space*: they adjust internal states dynamically to lower errors rather than relying solely on weight updates. This enables faster inference but at a cost—when faced with novel statistics, the past long-horizon observations make RNNs hallucinate context switches, imposing assumptions learned during training onto new situations.

We reasoned that an architecture that appropriately combines learning in the weight and activity space may be able to address the shortcomings of conventional RNNs. Accordingly, we developed NeuraGEM, a minimal neural architecture comprised of two interacting substrates, both updated to reduce output error but at distinct timescales. The difference in timescales results in a fast, decaying substrate that tracks rapid changes in the environment in activity space (*Z*), and a slow, stable substrate that consolidates accumulated knowledge in synaptic weights (*W*)(fig. 3A).

**Fig. 3.**
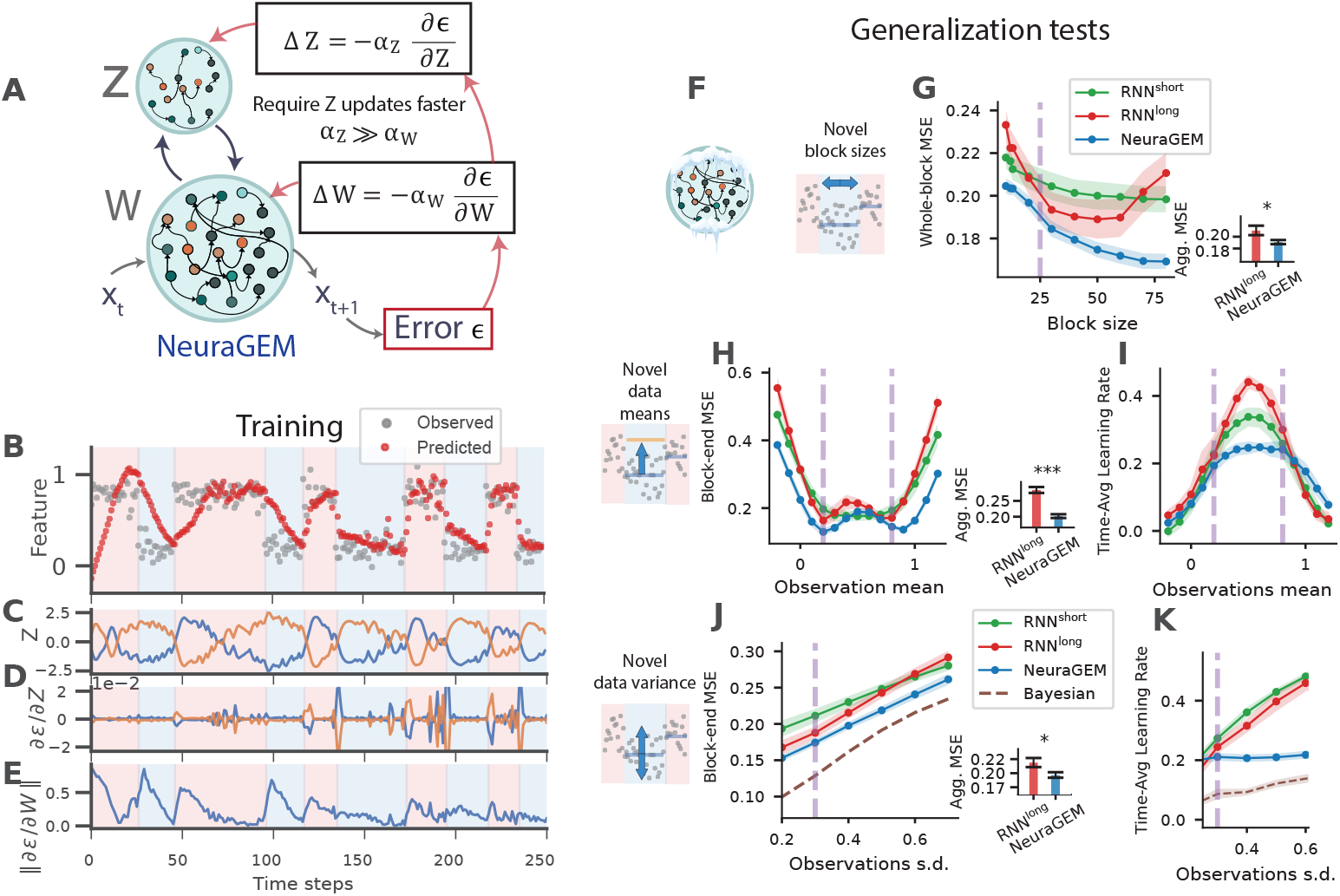
NeuraGEM discovers latent structure and improves generalization. **A**) Schematic of NeuraGEM with two interacting neural substrates, *Z* and *W* . The neural network *W*, modulated by *Z*, generates a prediction of the next sensory input and incurs a prediction error *ϵ*. Error is propagated back to both *Z* and *W*, which are updated to reduce output error, with *Z* updated at a faster rate. For this simulation, *Z* comprised two mutually inhibitory units that multiplicatively gate RNN_*W*_ . We used Adam optimizer with parameters: *α*_*Z*_ = 0.5, *α*_*W*_ = 0.001, 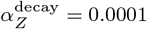 (we ignored 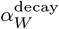). We used an input horizon of 5 time steps. **B**) Stimuli observed and NeuraGEM responses in the initial part of training (shown for Gaussian with s.d. = 0.1 for visual clarity). Background shading indicates the ground-truth context (0.2 or 0.8). **C**) Time series of *Z* activity during training. **D–E**) Gradients of the prediction error with respect to *Z* (**D**) and *W* (L1 norm; **E**). **(F)**: schematic indicating that weights were frozen during testing. **G**) Generalization to unseen block lengths. MSE across test block lengths for each model. Inset: Aggregate time-averaged MSE across block sizes (paired t-test, n = 20 independent model runs): RNN^long^ (mean ± s.e.m. 0.218 ± 0.008) vs NeuraGEM (0.191 ± 0.0035), *t*(19) = 2.34, *p* = 0.031. **H–I**) Generalization to unseen latent means. **(H)**: MSE for test means ranging from -0.2 to 1.2 (trained means indicated by dashed lines). Inset: Aggregate time-averaged MSE across mean values (paired t-test, n = 20): RNN^long^ (0.285 ± 0.0094) vs NeuraGEM (0.202 ± 0.006), *t*(19) = 8.0, *p <* 10^−4^. **(I)**: Corresponding behavioral learning rate across the same test range. **J–K**) Robustness to increased observation noise. **(J)**: MSE as observation standard deviation increases beyond the training level. Inset: Aggregate time-averaged MSE across noise levels (paired t-test, n = 20): RNN^long^ (0.219 ± 0.0081) vs NeuraGEM (0.197 ± 0.0049), *t*(19) = 2.50, *p* = 0.043. **(K)**: Behavioral learning rate under identical noise manipulations. The Bayesian model infers sample variance from the data. Shaded regions denote s.e.m. across 20 model seeds.

The principle of learning using two interacting substrates can be instantiated in various neural architectures (see Methods, section 3.1). To demonstrate how this mechanism learns context representations, we implemented a specific version of NeuraGEM and evaluated its performance on the one-dimensional context-switching prediction task introduced earlier, directly comparing its behavior to that of conventional RNNs.

The model comprises two interacting components (fig. 3A). The first is a recurrent neural network parameterized by a connectivity matrix *W* (RNN_*W*_) that predicts the next output. The second component consists of two mutually inhibitory units whose activity, *Z*, acts as a modulatory signal on RNN_*W*_ . The dynamics of *Z* are governed by two processes: an adaptive term that updates *Z* in the direction that reduces output error, and a leak term that drives *Z* to decay toward baseline. In general, the influence of *Z* on RNN_*W*_ can be multiplicative, additive, or follow more complex nonlinear forms. We used a multiplicative influence in all experiments (unless noted) with a static projection *Z* to RNN_*W*_ units.

Given an input *x*_*t*_, the model predicts the next sample

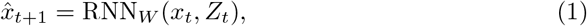

and incurs a prediction error

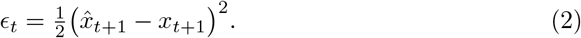

Both substrates are updated to minimize this error, but on distinct timescales:

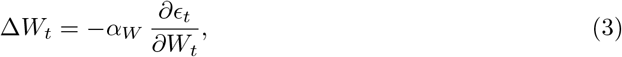

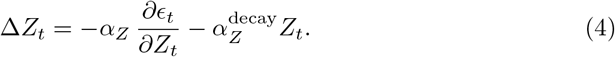

where *α*_*Z*_ is the update rate of *Z, α*_*W*_ is the update rate of *W*, and 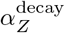 is the decay rate of *Z. W* and *Z* were updated using an Adam optimizer [29] with the values: *α*_*Z*_ = 0.5, *α*_*W*_ = 0.001, 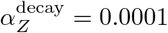.

While the output error *ϵ*_*t*_ is directly available to the network, adjusting *Z* in the correct direction is nontrivial because the output is produced by the recurrent network RNN_*W*_, which lies downstream of *Z*. Consequently, the influence of *Z* on the error is indirect. Updating *Z* therefore requires knowing how changes in its activity would alter the dynamics of RNN_*W*_ and, in turn, the resulting error. This dependency calls for an internal estimate of how *Z* affects the computations carried out by *W* —a mechanism analogous to internal forward models proposed in sensorimotor control [30, 31]. In our implementation, this sensitivity (*∂ϵ*_*t*_*/∂Z*_*t*_) is computed using standard backpropagation through time [32]. Later, we show that a similar computation can be achieved in a more biologically plausible way by introducing a learned feedback projection from RNN_*W*_ to *Z*, enabling local gradient estimation.

#### 1.2.1 Generalization performance of NeuraGEM

To evaluate how NeuraGEM learns and generalizes, we exposed the model to the same one-dimensional prediction task used for conventional RNNs (fig. 2). On each trial, the input *x*_*t*_ was sampled from a Gaussian distribution whose mean alternated covertly across blocks. As before, the model’s objective was to predict the next sample based on recent observations (h = 5 time steps), requiring it to infer the current latent context and adapt when the underlying distribution switched.

When trained on this task, NeuraGEM progressively separated the two latent distributions as it encountered block transitions (fig. 3B). Activity in *Z* came to represent each context uniquely (fig. 3C), while the error gradients with respect to *Z* increased and those with respect to *W* decreased—indicating a shift from slow synaptic adaptation to fast latent-state adjustment (fig. 3D,E). We compare these results to NeuraGEM with 10 *Z* units (showing similar dynamics), to NeuraGEM without *Z* decay (failing to converge), and RNN^short^ and RNN^long^ (see Supp. fig. 9).

Crucially, NeuraGEM outperformed both conventional RNNs and RNNs^long^ across all generalization tests (fig. 3F–K). It achieved a lower mean-squared error for both trained and novel block sizes (fig. 3G), maintained a lower error for unseen latent means where RNNs^long^ overfit (fig. 3H,I), and remained robust under increased observation noise (fig. 3J). The behavioral learning rate stayed low (fig. 3K), reflecting that random fluctuations in input did not systematically bias *Z* and were effectively integrated away.

### 1.3 NeuraGEM as a neural analogue of Expectation–Maximization

A useful way to understand how NeuraGEM clusters latent task states is through its correspondence to the Expectation–Maximization (EM) algorithm, a classical optimization method for discovering hidden structure in data. EM provides a conceptual template for how a system can infer latent categories while simultaneously refining the parameters that describe them. The algorithm alternates between two operations: (1) estimating which latent category each data point is assigned to (E-step) and (2) updating the model parameters based on those assignments (M-step) (fig. 4A). Online variants of EM have been used for time-series latent-variable models [25, 26, 33], including neural network–based formulations [34–36].

**Fig. 4.**
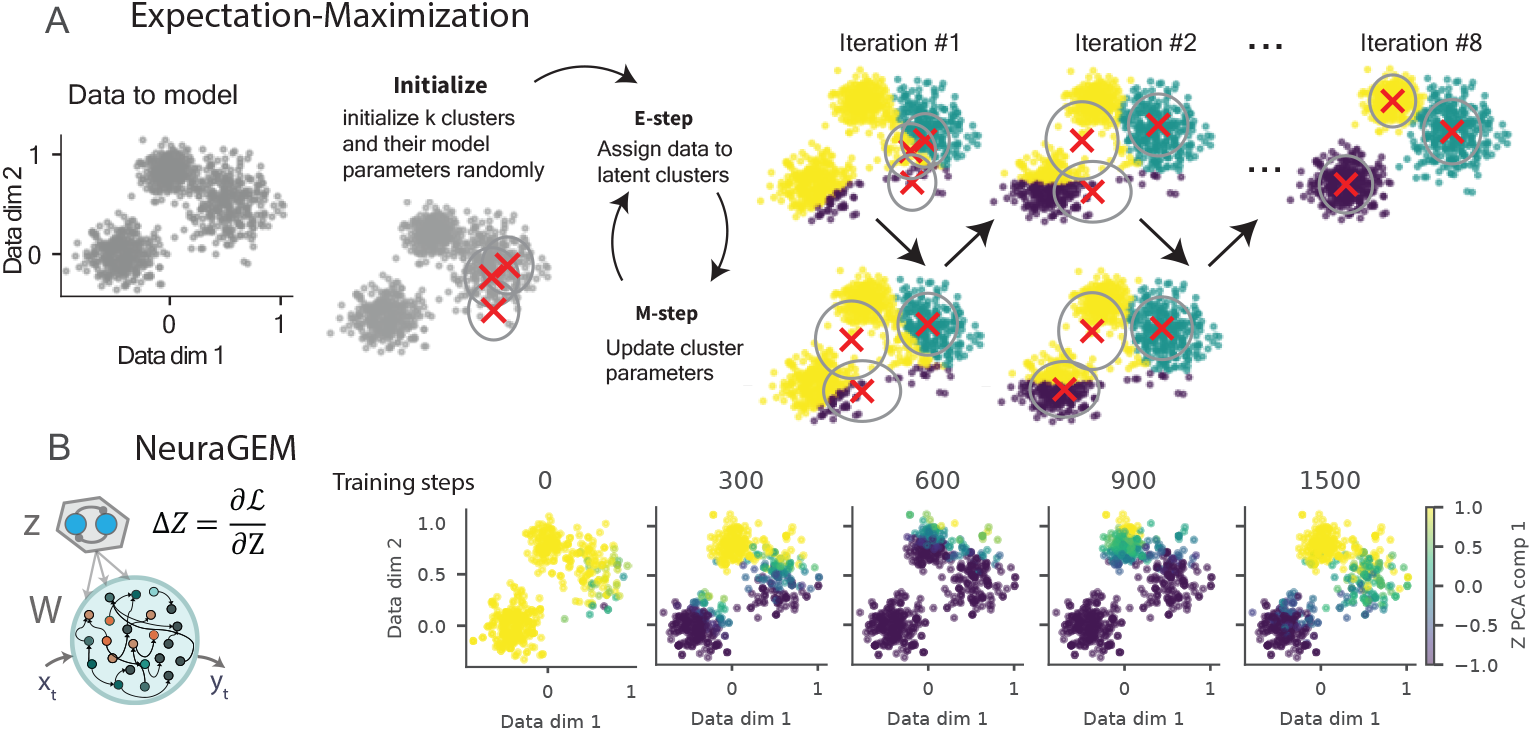
NeuraGEM functions similarly to the Expectation-Maximization algorithm. **A**) Schematic illustration of the Expectation–Maximization (EM) algorithm on 2D data drawn from three Gaussian clusters (left panel). Each cluster model is shown as a cross (cluster mean) and an ellipse (covariance). Points are colored according to their current cluster model assignment. The right panels show the iterative EM process: the E-step reassigns points to the nearest cluster model (color changes), and the M-step updates each model’s mean (cross) and covariance (ellipse). **B**) NeuraGEM analogue of cluster assignments. The model was trained on temporal data drawn from the same three Gaussian clusters presented in blocks. To obtain an EM-like “assignment,” the model predicted each observation and *Z* was optimized to lower error, while holding W fixed. The resulting *Z* timeseries was projected onto the first principal component, shown as a continuous color gradient. The sub-panels display how these projected values evolved over training.

NeuraGEM can be derived from the same objective that underlies EM: maximizing the marginal likelihood of observations under a model with latent variables (derivation in supplement 4.1). NeuraGEM replaces the E-step and M-step operations with gradient-based updates driven by prediction error (Equations 4, 3). Updates to *Z* to lower prediction errors correspond to an E-step that searches for the most likely assignment of *Z*, whereas the slower weight update (Equation 3) optimizes the parameters *W* for the current inferred *Z*, corresponding to the M-step.

A critical ingredient is the separation of timescales *α*_*Z*_ ≫ *α*_*W*_ . *Z* equilibrates faster than *W* ; thus, as *W* is updated, *Z* is close to its optimum for the current *W*, an important assumption in EM. The decay term in the *Z* update pulls latent activity back toward a baseline in the absence of supporting evidence. This prevents runaway activity and ensures that new contexts are only maintained when they consistently reduce prediction error, promoting a stable and parsimonious latent structure (Supp. fig. 9C–E, see Supplement section 4.1).

NeuraGEM clusters evolve throughout training with dynamics comparable to EM. For a qualitative comparison, we trained NeuraGEM on the same 2D data drawn from three clusters, but presented the data in blocks (fig. 4). The *Z* population, implemented as two mutually inhibitory units in soft competition, rapidly aligned with the cluster most consistent with the current input (fig. 4B). The alignment between *Z* activity and ground-truth clusters improved gradually throughout training, similar to EM dynamics. Although the two units are mirror-symmetric, their graded population activity can represent multiple latent clusters through distributed coding.

### 1.4 NeuraGEM discovers latent structure across multiple timescales

Natural environments often contain multiple latent causes that evolve on different timescales—some change rapidly, while others drift over extended periods. A single inference process cannot handle this diversity: a system tuned for fast changes will misinterpret slow structure as noise, whereas a system tuned for slow changes will overlook rapid transitions.

We reasoned that intrinsic differences in how *Z* units accumulate and forget might enable NeuraGEM to make appropriate inferences when latent contexts span multiple timescales. Accordingly, we instantiated a model with two Z populations that differed only in their update and decay constants. The “fast” population, *Z*_fast_, had larger update and decay rates, allowing it to react strongly but transiently to errors. The “slow” population, *Z*_slow_, integrated errors more gradually and retained information more effectively (fig. 5A).

**Fig. 5.**
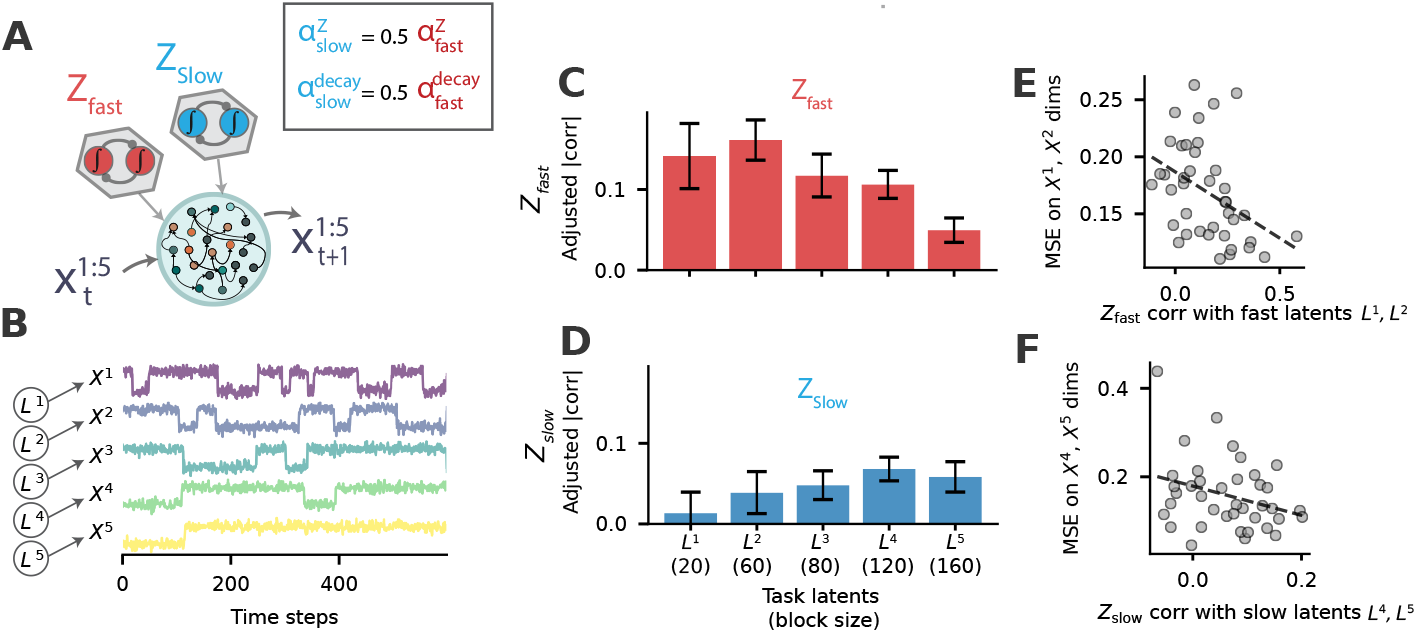
Small differences in *Z* dynamics modulate sensitivity to environment timescales. **A)** Schematic of NeuraGEM extended with two latent populations: a fast population (*Z*_fast_) with larger update and decay rates (*α*_*Z*_ = 1.0, 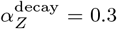) and a slow population (*Z*_slow_) with smaller rates (*α*_*Z*_ = 0.5, 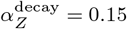). Note that this simulation uses the Stochastic Gradient Descent optimizer (SGD) which required higher hyperparameter values compared to Adam optimizer used before. The Adam optimizer offers robustness by dynamically adapting learning rates, which obscures the time scale differences examined here. **B**) Five-dimensional input stream generated by corresponding five latent variables that switch at distinct timescales. Each latent switches between 0.2 and 0.8 and accordingly sets the Gaussian mean value to generate observations in the corresponding input dimension. **C**) Post-training correlation between *Z*_fast_ activity and each latent variable. **D**) Correlation between *Z*_slow_ activity and the latent variables. Bars show mean adjusted correlations across 10 runs; error strips denote SEM. **E**) Scatter plot of Z_fast_ correlation with the first two latent variables (*L*^1^, *L*^2^, block sizes 20 and 60) and the prediction MSE on the corresponding input dimensions (X^1^ and X^2^). Each point denotes one observation formed by a run (seed) *×* latent dimension. Dashed line shows the least-squares fit. Pearson correlation (two-tailed): *r* = −0.407, *p* = 0.0092, *df* = 38, *n* = 40 observations (*R*^2^ = 0.165). **F**) Same as in (**E**) but for Z_slow_ correlation with the slower latent variables (*L*^4^, *L*^5^, block sizes 120 and 160) and prediction MSE along the corresponding inputs (X^4^ and X^5^). Pearson correlation (two-tailed): *r* = −0.305, *p* = 0.0591, *df* = 38, *n* = 40 observations (*R*^2^ = 0.093). Cross-tests with Z_fast_ correlation with the slower latent variables and Z_slow_ correlation with the faster latent variables show smaller *R*^2^ values (Supp. fig. 10).

We evaluated this architecture in a task designed to contain multiple latent processes operating at different timescales. The input stream consisted of five-dimensional observations with five independent latent variables, each generating one dimension of the input (fig. 5B). Crucially, each latent variable switched its mean at a distinct rate, ranging from rapidly changing blocks of 20 samples to slowly changing blocks of 160 samples.

After training, the two populations exhibited clear and complementary specialization. *Z*_fast_ aligned most strongly with the latent dimensions that changed quickly, while *Z*_slow_ selectively tracked the latent variables that evolved over the longest timescales (fig. 5C,D). This alignment with the underlying latent variables improved task performance. The degree to which *Z* tracked a latent variable was associated with lower prediction errors in the corresponding observation dimension (fig. 5E,F). Importantly, this specialization was not imposed by the architecture or by supervision; it emerged spontaneously from the slight differences in update and decay dynamics. This finding suggests that NeuraGEM–when equipped with small amounts of heterogeneity– can handle relatively complex contextual inference problems [37, 38].

### 1.5 Modeling sequence learning in humans

We next asked whether NeuraGEM can account for certain counterintuitive features of human sequence learning. We turned to a recent study in which participants predicted outcomes in short “stories” generated by one of two hidden Markov chains [39]. Each story contained a transition that uniquely identified the latent context, followed by a transition whose outcomes were randomized and shared across contexts (fig. 6A). Although the diagnostic cue was perfectly informative of the current context, human performance depended strongly on the curriculum with which the contexts were experienced.

**Fig. 6.**
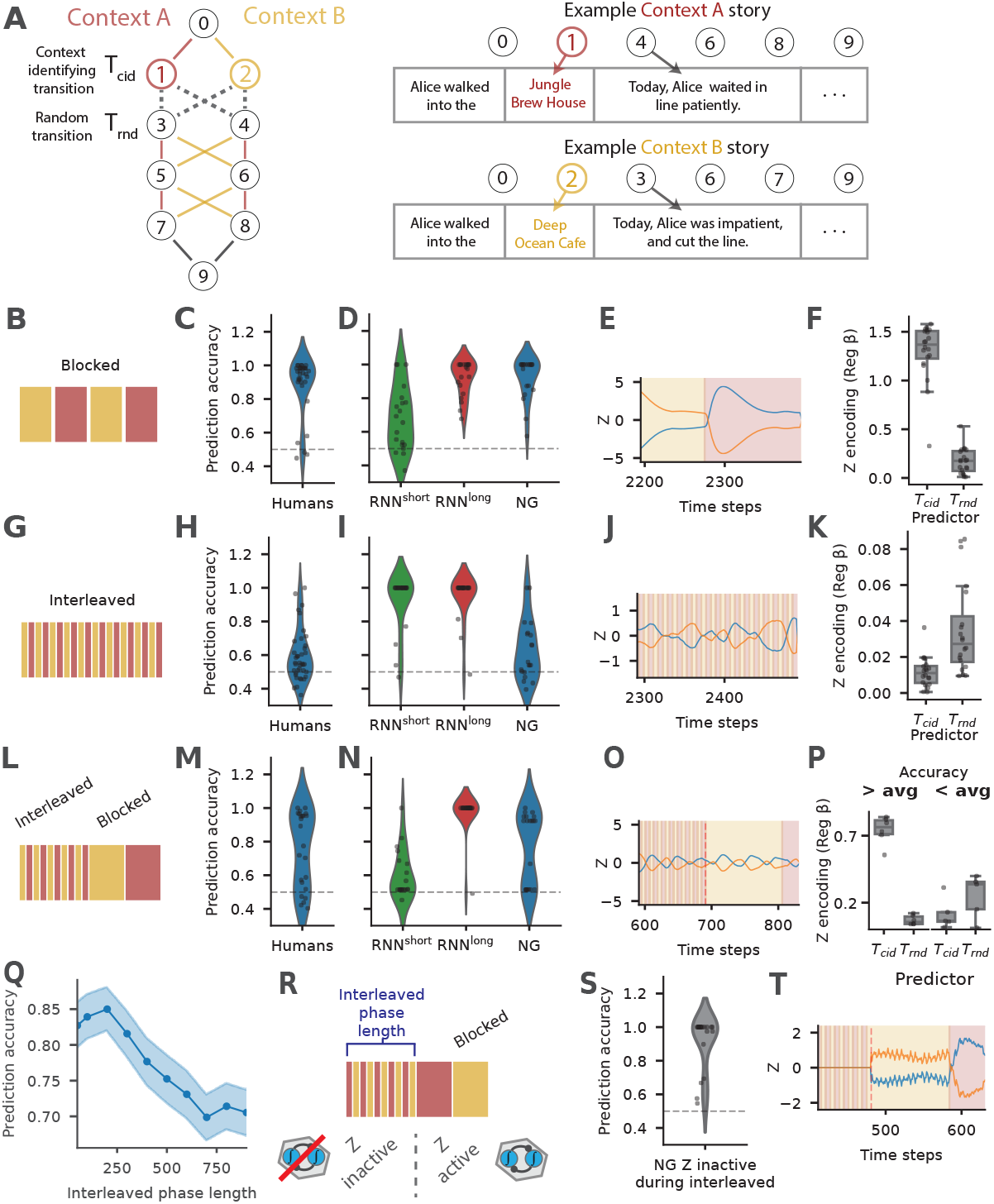
Behavior and model responses under different training curricula in a sequence learning task. **A**) Structure of the sequence learning task. Left: two Markov chains define the latent contexts, with solid edges indicating deterministic transitions (including the context-identifying transition *T*_*cid*_) and dotted edges indicating random transitions (*T*_*rnd*_). Right: example story outcomes corresponding to specific states. **B, G, L**) Training curricula. Blocked training (**B**) alternates contexts in long segments; interleaved training (**G**) alternates contexts every story; mixed training (**L**) begins with an interleaved phase followed by a blocked phase. **C, H, M**) Prediction accuracy for human participants under blocked (**C**), interleaved (**H**), and mixed (**M**) training. Adapted from [39] with permission. **D, I, N**) Prediction accuracy for RNN^short^, RNN^long^, and NeuraGEM under the same three curricula: blocked (**D**), interleaved (**I**), and mixed (**N**). For both human participants and the models, prediction accuracy following each of the training curricula was tested using a random curriculum (see Methods section 3.5). **E, J, O**) Time series of NeuraGEM latent activity *Z*, with shaded backgrounds indicating the underlying context. **F, K, P**) Regression coefficients (*β*) relating *Z* activity to *T*_*cid*_ and *T*_*rnd*_ for the three curricula. In **P**, runs are grouped by above- and below-average final accuracy. OLS linear regression of (Z) on transition predictors (including (*T*_*cid*_) and (*T*_*rnd*_)); absolute regression coefficients reported. Boxplots show median and IQR; dots denote individual runs (n = 20). **Q**) Prediction accuracy as a function of interleaved-phase duration (x-axis) for NeuraGEM. Shaded regions denote SEM across 20 runs. **R-T**) Manipulation in which *Z* updates are disabled during the interleaved phase. (**R**): shows the mixed curriculum but with Z inactive during interleaved phase and active during blocked phase. (**S**): shows end prediction accuracy under this condition; (**T**): shows the corresponding *Z* activity around the transition from interleaved to blocked training.

Under **blocked training**, where the same context persisted for many consecutive stories, participants quickly learned the generative structure (fig. 6C). NeuraGEM showed the same pattern, using only a short input horizon (h=6 time steps, reflecting evidence that humans flush the contents of working memory at event boundaries [40]). The model reached human-level accuracy, and its fast variable *Z* reliably tracked the underlying context (fig. 6D-F, h=10 and 20 had qualitatively similar results). A standard RNN failed in this regime, and RNN^long^ trained with long horizons succeeded only with substantially long training input sequences (h= 200 time steps) (fig. 6D).

Under **interleaved training**, where the generating context alternated on every story, human learners struggled (fig. 6H). NeuraGEM struggled in the same manner. Rather than aligning with the true context, *Z* locked onto the random transition, which happened to show weak incidental structure (fig. 6I-K). Both conventional RNNs^short^ and RNNs^long^ performed well in this setting, missing the human failure entirely (fig. 6I).

Previous modeling efforts attributed the learning inefficiency for interleaved curricula to mismatches between task demands and internal context dynamics [41–43]. While we also observe that *Z* has a temporal sensitivity, our results indicate an additional mechanism: multiple features compete to entrain *Z*. The random transitions *T*_*rnd*_ contain incidental temporal structure that competes with the context transitions *T*_*cid*_, leading *Z* to encode one at the expense of the other (their encodings are anti-correlated, Supp. fig. 11D). This makes a counterintuitive and testable prediction: matching the temporal patterns of the two transitions (by making the context transitions random instead of interleaved) should rescue a portion of interleaved training runs (Supp. fig. 11).

A **mixed curriculum** produced a more revealing signature. When interleaved exposure preceded blocked exposure, human performance split: some learners recovered once the structure stabilized, while others remained impaired [39]. NeuraGEM reproduced this bimodality (fig. 6L-N). In successful runs, *Z* shifted to the diagnostic cue once blocked structure appeared; in failed runs, early mis-clustering persisted, and the model continued to track the irrelevant transition (fig. 6O,P). The depth of this misalignment during the interleaved phase predicted whether recovery would occur (Supp. fig. 12).

The model makes two additional experimental predictions. First, impairment due to interleaved training should scale with the duration of interleaved exposure: longer interleaved phase increases the probability of persistent failure (fig. 6Q). Second, preventing context discovery during interleaved training should rescue later blocked training. Consistent with this, disabling *Z* updates during the interleaved phase fully restored performance under subsequent blocked training (fig. 6R-T).

Taken together, these results show that NeuraGEM captures both the successes and the failures of human learners in this task. The model’s behavior reflects a simple principle: with limited evidence, fast latent updates can stabilize around the wrong structure, and early misalignment can persist even when clearer information becomes available. This mechanism offers a compact account of why curriculum matters and accounts for the observation that humans sometimes fail to recover from early misleading experience.

### 1.6 Dissecting dynamical mechanisms in NeuraGEM

We next used the earlier one-dimensional prediction task to examine how NeuraGEM’s architecture shapes its internal representations and dynamics. Our aim was to understand the computational structure and how it might differ from that in conventional RNNs. To do so, we analyzed three aspects of the learned networks: (i) how each model responds to the statistical structure of the inputs, (ii) how context information is encoded in population activity, and (iii) how each model’s vector field shapes its corresponding latent dynamics.

We first analyzed the dependency of each model’s output on the underlying statistics of the observed samples (fig. 2A). Models were trained on blocks of Gaussian samples with a mean alternating between 0.2 and 0.8 but were tested on a range of other mean values. The RNN^short^ behaved like a local estimator: it regressed strongly toward the mean of the most recently experienced context (fig. 7A). The RNN^long^ developed two strong attractors at the trained means and showed repulsion for intermediate values, producing strong training-dependent biases (7B). NeuraGEM showed a qualitatively different pattern: *Z* values organize the input-response relationship such that prediction errors directly move the system onto the appropriate response curve. As a result, NeuraGEM is able to rapidly and flexibly adapt to novel mean values (fig. 7C).

**Fig. 7.**
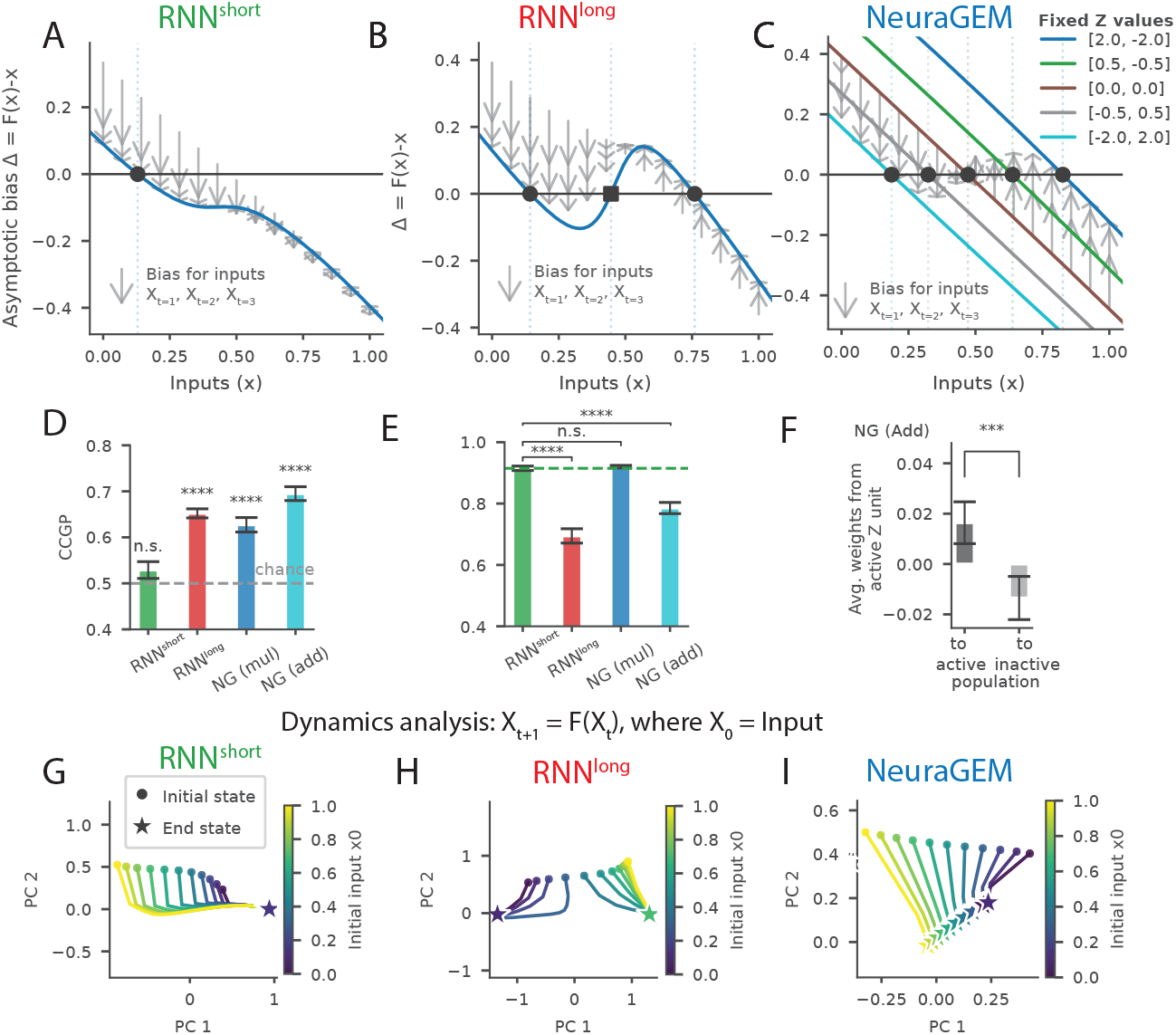
Neural analyses of recurrent models and NeuraGEM. **A–C**) Response bias (Δ = output – input) as a function of input probe value. Solid curves show the asymptotic bias towards the end of a block. Arrows denote the change in output bias in response to the first three input samples *X*_1:3_. Zero crossings indicate fixed points toward which responses regress (circle markers) or from which they are repelled (square markers). (A) RNN^short^ exhibits a single regression point determined by observations seen in the last block of training. (B) RNN^long^ exhibits two stable attractors corresponding to the trained contexts, with repulsion for intermediate values. (C) NeuraGEM bias curves plotted for several fixed values of *Z*, illustrating strong modulation of model behavior by the Z activity. Arrows show bias following the first three input samples, during which rapid updates of *Z* move the system toward the response curve most consistent with the input values. **D**) Cross-condition generalization performance (CCGP) for decoding context from hidden-unit activity (see Methods). Chance level is 0.5 (dashed). Significance relative to chance was assessed using two-sided one-sample *t*-tests across seeds: RNN *t* = 1.58, *p* = 1.30 *×* 10^−1^ (n.s.); RNN^long^ *t* = 15.52, *p* = 3.02 *×* 10^−12^ (****); NeuraGEM (multiplicative) *t* = 8.07, *p* = 1.47*×*10^−7^ (****); NeuraGEM (additive) *t* = 12.94, *p* = 7.20*×*10^−11^ (****). **E**) Cosine similarity of hidden population activity across successive context blocks. Lower values indicate greater separation between context-specific population activity. Each model was compared to the short-horizon RNN baseline using two-sided paired *t*-tests across seeds: RNN^long^ vs RNN^short^ *t* = 9.02, *p* = 5.91 *×* 10^−9^ (****); NeuraGEM (multiplicative) vs RNN^short^ *t* = −0.63, *p* = 5.31 *×* 10^−1^ (n.s.); NeuraGEM (additive) vs RNN^short^ *t* = 6.59, *p* = 6.72 *×* 10^−7^ (****). **F**) Learned structure in projections from the fast neural substrate *Z* to recurrent units in the additive NeuraGEM variant. Hidden units were grouped by context preference based on mean activity. For each seed, weights from the most active *Z* component were averaged separately over units preferring each context. A two-sided paired *t*-test across seeds revealed a reliable asymmetry: *w*(*z*_1_) vs *w*(*z*_2_), *t* = 4.14, *p* = 5.56 *×* 10^−4^ (***). **G–I**) Autonomous dynamics visualized by projecting population activity into state space. Models were initialized with different inputs and then evolved without external drive by recursively feeding their output as the next input. (G) RNN^short^ trajectories converge to a single fixed point. (H) RNN^long^ trajectories converge to two fixed points. (I) NeuraGEM trajectories form a continuum of states parametrized by *Z*. Error bars denote s.e.m. across 20 seeds. Additional details of metrics and analyses are provided in Methods.

We next examined how each model encodes context within its population activity using cross-condition generalization performance (CCGP) and the cosine similarity between units activated for each context. High CCGP indicates that the neural state cleanly separates the two contexts along a low-dimensional axis invariant to noise or stimulus level [9]. The RNN^short^ showed chance-level CCGP, accompanied by the activation of an overlapping population of units in each context (fig. 7D, E). By contrast, the RNN^long^ achieved high CCGP and recruited relatively separated neural populations for each context (fig. 7D, E).

For NeuraGEM, we examined two different instantiations: one in which *Z* influenced *W* multiplicatively (mul) and another in which the effect was additive (add). Both variants reached high CCGP values (fig. 7D), indicating robust stimulus-invariant context representations. However, the multiplicative variant mostly activated the same population of units in each context (despite gating prior to recurrent weights), while the additive variant activated separate populations (fig. 7E). This is supported by the structure in the Z to RNN_*W*_ weights in the additive variant, where each Z unit preferentially activated one of the two functional neural populations (fig. 7F), consistent with Z activity acting as a control variable, switching neural pathways to direct computations [44].

Finally, we characterized the latent dynamics of each model by evaluating its recurrent behavior as a trajectory on a vector field. To do so, we initialized each network with a sensory input and then allowed it to evolve recursively without further external drive. RNN^short^ converged to a single attractor regardless of initialization (fig. 7G). RNN^long^ settled into one of two attractors corresponding to the trained means (fig. 7H). In contrast, NeuraGEM converged to a continuum of states parameterized by *Z*. This continuum forms a line attractor, providing a dynamical substrate for smoothly adjusting internal states to represent new or intermediate contexts [45] (fig. 7I).

### 1.7 NeuraGEM with a learned feedback mechanism

Updating *Z* based on error is nontrivial because *Z* influences the output only indirectly through a recurrent network, RNN_*W*_ . In the results presented thus far, we used standard backpropagation through time [32] to adjust *Z* even in the test phase. Although convenient, this requires differentiating through the RNN_*W*_ weights and activity dynamics, which is neither efficient nor biologically plausible.

To address this limitation, we need a compact mechanism through which errors could more directly alter *Z*. We reasoned that this may be achieved by a separate feedback network RNN_*f*_ that transforms output errors into appropriate *Z* updates [46]. Accordingly, we trained an additional network to serve as a learned feedback pathway that computes *Z* updates directly from the output error (See Methods section 3.7). This feedback pathway operates in parallel with the forward network, preserving performance and enabling context switching (fig. 8).

**Fig. 8.**
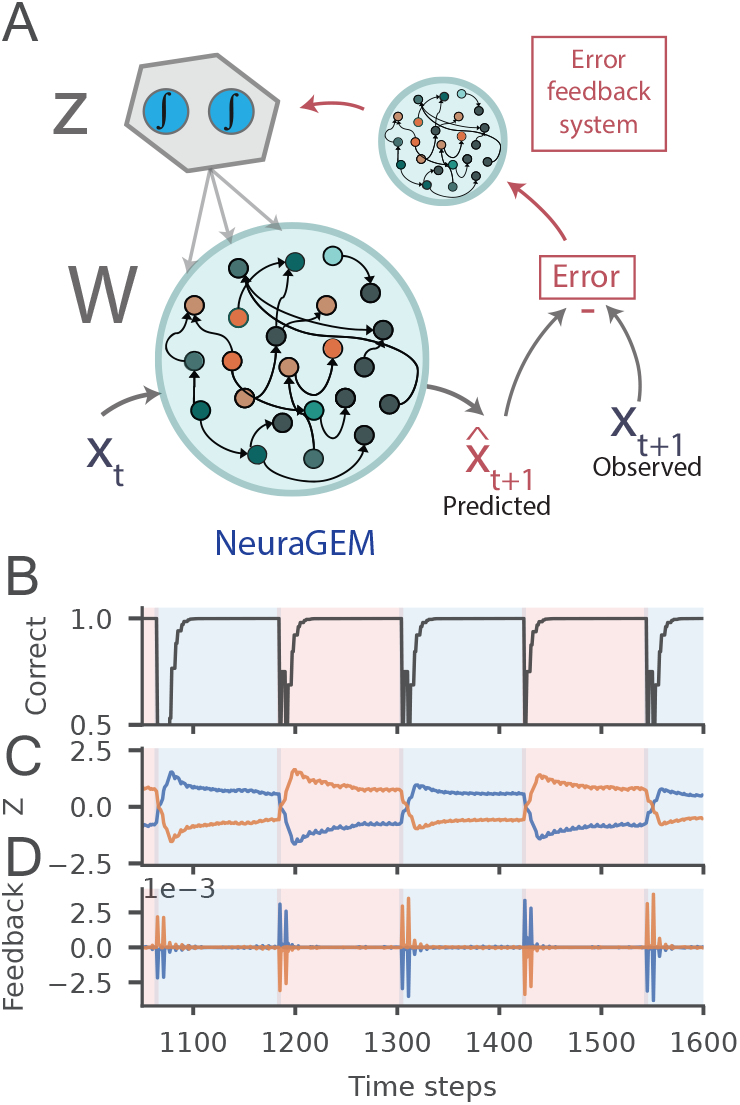
A learned feedback pathway switches context without backpropagation. **A**) Schematic. The forward network RNN_*W*_ predicts 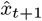 and yields an instantaneous residual 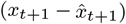. A separate feedback network RNN_*f*_ takes this residual together with *Z*_*t*_ and outputs a gradient Δ*Z*_*t*_ used to update *Z* with decay 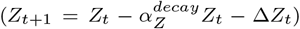. **B**) Prediction accuracy after training on the sequence learning task (section 1.5) **C**) *Z*_*t*_ aligns with block identity, switching rapidly at boundaries. **D**) Predicted gradients Δ*Z*_*t*_ (output of RNN_*f*_) exhibit sharp, switch-locked spikes at block transitions, reflecting boundary-sensitive error bursts.

**Fig. 9.**
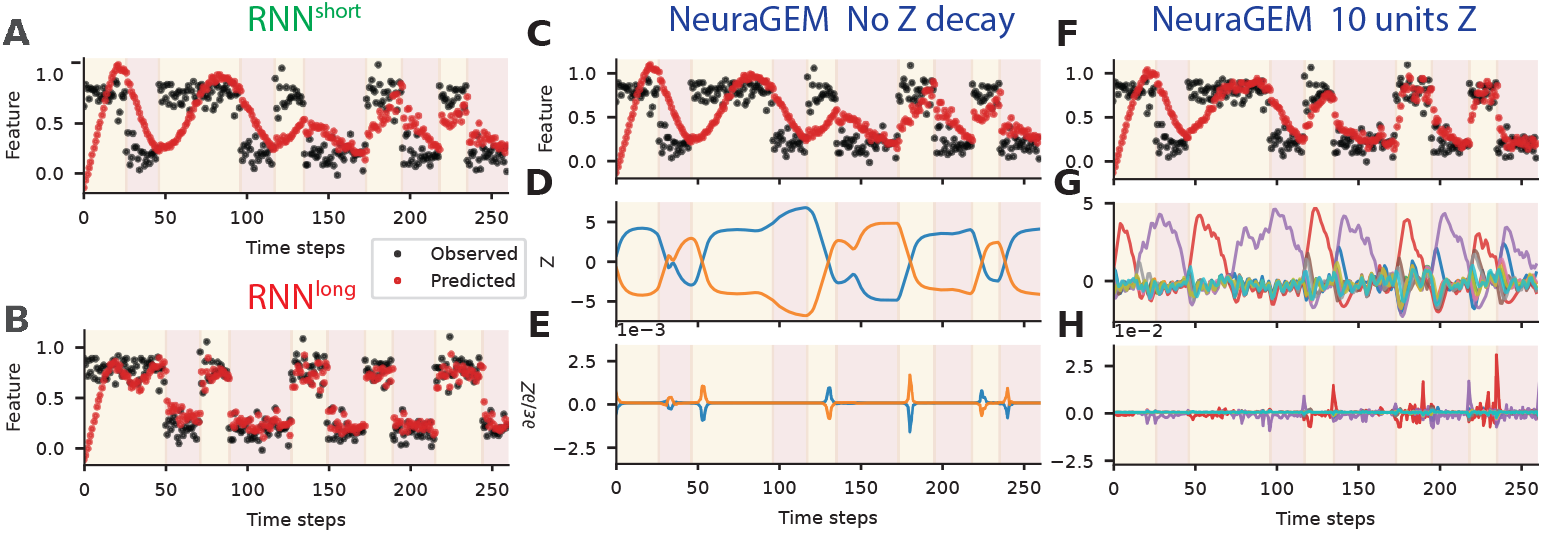
Training of RNNs and RNN^long^s. A) Observations and RNN predictions from the initial training stretch of an example short-horizon (h=5) RNN, and B) a long-horizon (h=50) RNN. Note that long-horizon RNN consumes the first 50 time steps in its first training time step prior to any weight updates and those time points were excluded. C-E) NeuraGEM run on the same observations but with 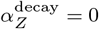 F-G) NeuraGEM run on the same observations but with 10 units in *Z*.

**Fig. 10.**
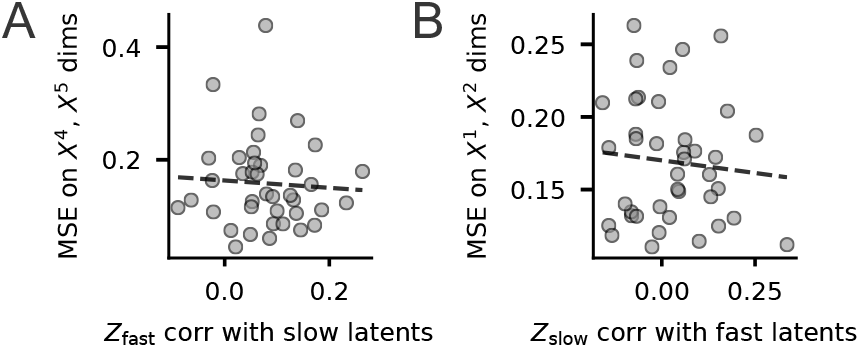
Cross-tests for Z_fast_ and Z_slow_ correlation with prediction MSE on opposite input dimensions. **A**) Scatter plot of Z_fast_ correlation with the slower latent variables (block sizes 120 and 160) and the prediction MSE on the corresponding input dimensions (X^4^ and X^5^). Pearson correlation (two-tailed): r=-0.063, p=0.7033, df=37, n=39, *R*^2^=0.004. **B**) Same as in (**E**) but for Z_slow_ correlation with the first two latent variables (block sizes 20 and 60) and prediction MSE along the corresponding inputs (X^1^ and X^2^). r=-0.095, p=0.5612, df=38, n=40, *R*^2^=0.009.

**Fig. 11.**
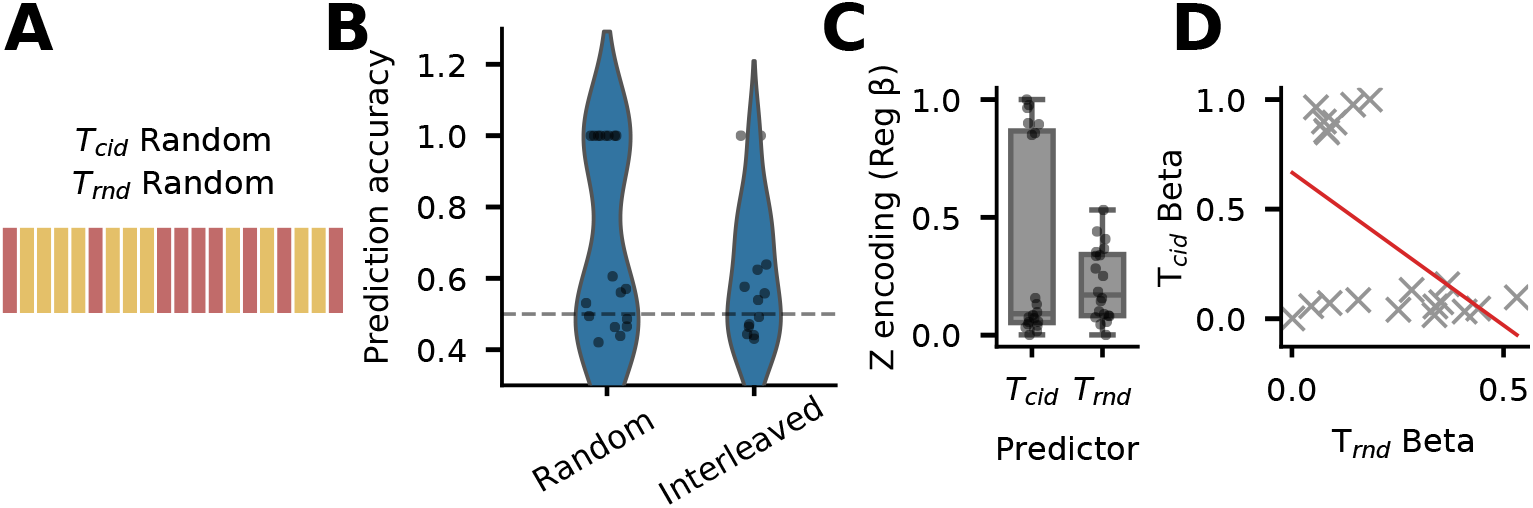
Competing temporal structures biases latent-state encoding in NeuraGEM. Under an interleaved curriculum, NeuraGEM preferentially tracks the random transition (*T*_rnd_), resulting in poor prediction accuracy. We hypothesized that this failure arises because incidental temporal structure in *T*_rnd_ resonates with the intrinsic dynamics of *Z*. To test this, we designed curricula in which *T*_cid_ and *T*_rnd_ were matched in their temporal statistics. **A**) Stories were generated and randomly shuffled such that both the context-identifying transition (*T*_cid_) and the random transition (*T*_rnd_) appeared in random order. **B**) End-of-training accuracy when the context-identifying transition was random, compared to when the context-identifying transition was interleaved. **C**) Regression coefficients relating *Z* activity to outcomes of *T*_cid_ and *T*_rnd_. **D**) Scatter plot of regression coefficients reveals a trade-off: stronger alignment of *Z* with one transition is associated with weaker alignment with the other. (OLS regression, *n* = 20, slope = −0.342, *r* = −0.47, *t*(18) = −2.24, two-sided *p* = 0.038).

**Fig. 12.**
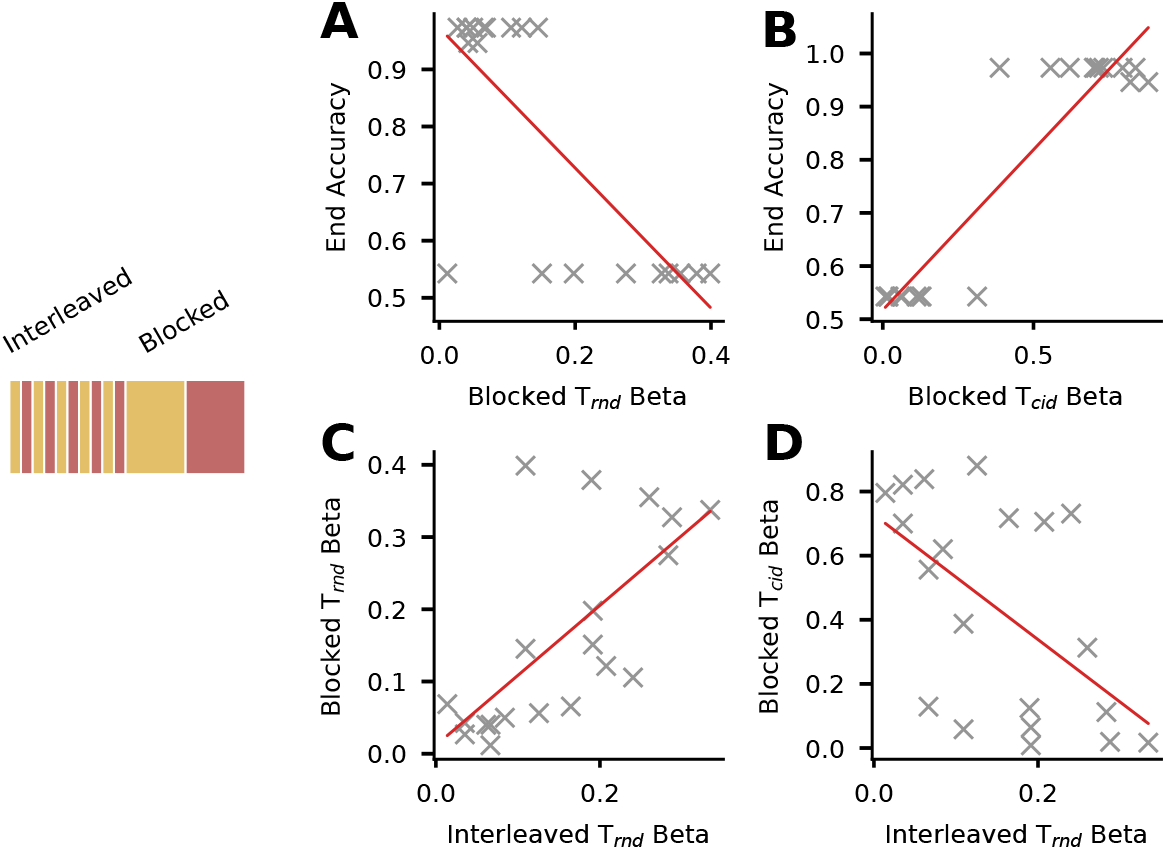
Persistence of *Z* representations across curricula predicts final performance. To quantify what the fast neural substrate *Z* encoded at different stages of training, we regressed *Z*_*t*_ activity onto story transition predictors and extracted regression coefficients (betas) for the contextidentifying transition (*T*_cid_) and the random transition (*T*_rnd_). Betas were computed over the entire interleaved phase and over the entire blocked phase, and related to end-of-training prediction accuracy. **A**) Relationship between the *T*_rnd_ beta during the blocked phase and final prediction accuracy. **B)** Relationship between the *T*_cid_ beta during the blocked phase and final prediction accuracy. **C**) Correlation between *T*_rnd_ betas estimated during the interleaved phase and during the blocked phase, indicating persistence of *Z* encoding across curricula. **D**) Relationship between the *T*_rnd_ beta during the interleaved phase and the *T*_cid_ beta during the blocked phase, assessing whether early misalignment predicts later context encoding. **Regression statistics**. Ordinary least-squares (OLS) linear regression with two-sided *t*-tests on the slope (*n* = 20 model runs). Blocked *T*_*rnd*_ beta vs end accuracy (OLS linear regression, two-sided t-test on slope, n = 20): slope = -1.298, *r* = −0.72, *t*(18) = −4.44, *p <* 10^−3^. Blocked *T*_cid_ beta vs end accuracy (OLS linear regression, two-sided t-test on slope, n = 20): slope = 0.626, *r* = 0.85, *t*(18) = 6.84, *p <* 10^−4^. Interleaved *T*_*rnd*_ vs blocked *T*_*rnd*_ (OLS linear regression, two-sided t-test on slope, *n* = 20): slope = 0.810, *r* = 0.65, *t*(18) = 3.59, *p* = 0.002. Interleaved *T*_*rnd*_ vs blocked *T*_*cid*_ (OLS linear regression, two-sided t-test on slope, *n* = 20): *slope* = −1.639, *r* = −0.54, *t*(18) = −2.69, *p* = 0.015.

RNN_*f*_ learned to produce effective Δ*Z*_*t*_ transients precisely at latent context transitions, even though no explicit switch cue was provided (fig. 8D). These events were driven purely by prediction errors and selectively marked covert changes in context. The capacity to transform performance errors into activity updates mirrors post-error changes in firing rates in frontal cortical areas [12, 16].

## 2 Discussion

Flexible behavior requires the ability to infer latent structure from ongoing experience and to update these inferences when the environment changes. Here, we introduced NeuraGEM, a simple two-timescale neural architecture that discovers and tracks latent task states through the interaction of fast, error-driven activity, and slow synaptic plasticity. This division of labor implements an online analogue of the Expectation–Maximization algorithm: fast updates assign recent observations to putative latent states, while slow plasticity stabilizes the corresponding task modules. Through this mechanism, NeuraGEM captures core behavioral, computational, and neurophysiological features of latent-state inference.

A central insight from our work is that rapid context inference does not require deep unrolled histories, large training sets, or explicit supervision. Conventional recurrent networks either update too slowly (when relying on synaptic changes) or overfit to long-horizon training statistics (when leveraging meta-learning dynamics). By contrast, NeuraGEM separates two roles: a fast process that adjusts internal activity in response to moment-to-moment prediction errors, and a slow process that learns stable structure across time. This interaction allows the model to detect changes in latent context as soon as they occur, while also preserving the representations needed for long-term generalization.

A striking departure of NeuraGEM from optimal behavior captured a counterintuitive feature of human contextual learning. In a sequential prediction task, the model detected covert structure rapidly under blocked exposure but exhibited persistent misclustering after early interleaved exposure—mirroring the paradoxical curriculum effects observed in human learners. The model also predicted the conditions under which recovery should occur and identified manipulations that selectively prevent such misclustering. These results suggest that failures to recover from misleading early structure may reflect the system becoming trapped in a local latent-state minimum.

At the neural level, NeuraGEM exhibits representational geometries and dynamics consistent with observations in frontal circuits. The model forms low-dimensional context representations analogous to those reported in prefrontal cortex [9], and shows sharp transient responses at latent-state transitions reminiscent of error signals in the anterior cingulate cortex [12, 16]. Its intrinsic dynamics organize into a line attractor parameterized by the latent state, providing a dynamical substrate for smoothly tracking contextual changes [45, 47–49].

Neural network models gated by context representations provide a compelling account of flexible control, in which context signals simplify learning and promote generalization. Many such models, however, either assume the availability of context representations or rely on task-specific gating mechanisms, whose applicability across timescales and learning regimes remains unclear [21, 47, 50–54]. Even explicitly modular approaches, such as mixture-of-experts architectures, require a prescribed decomposition into experts to ensure stable specialization [55–58].

By contrast, NeuraGEM leverages the temporal structure of experience to induce task modules through learning, without predefining their number, form, or assignment. In this respect, NeuraGEM is computationally related to work in cognitive science showing that humans infer latent task structure and organize learning accordingly [1, 7, 59–62], while providing a simple neural mechanism grounded in interacting error-driven processes.

Previous work on error-minimizing circuits—most notably predictive-coding and its temporal variants—frames adaptation as continuous inference over hidden causes, well suited to gradual changes in sensory statistics [63–66]. NeuraGEM complements this view by showing how a separation of timescales can convert error signals into abrupt, discrete updates of latent state.

Our work also highlights several avenues for future research. First, how can this architecture scale to richer and more numerous latent structures? Natural tasks often involve hierarchies of contexts and partially overlapping causes. Extending NeuraGEM to such domains will likely require distributed or hierarchical implementations of the fast component, enabling multiple latent states to be inferred and coordinated across levels of abstraction.

Second, a critical open question concerns the circuit mechanisms that could implement fast latent updates without non-local gradient transport. Although we demonstrated that a learned feedback pathway can approximate the necessary update direction after learning, future work should examine whether and how such feedback circuits support learning through error-to-state transformations.

Third, the model generates concrete predictions that can be validated experimentally. For example, it predicts that context switches are accompanied by rapid, transient reorganization in activity space, followed by slower synaptic consolidation. This separation of timescales implies measurable signatures: early boundary-locked activity transients, coordinated changes in effective connectivity, and vector-field dynamics that reorganize around attractor structures aligned with environmental statistics. Testing these predictions will require joint measurements of fast neural activity, feedback interactions, and slower plasticity-related changes during learning.

In sum, NeuraGEM offers a compact and testable mechanism by which neural circuits can rapidly infer hidden task states to support adaptive behavior and generalization.

## 3 Methods

Code to reproduce all figures and experiments in PyTorch is available at: https://github.com/hummosa/neuragem. All reported results use 20 random seeds unless noted otherwise; summary statistics are reported as mean ± s.e.m. across seeds.

### 3.1 NeuraGEM: general formulation

NeuraGEM is a two-timescale neural architecture composed of a fast neural substrate *Z* and a slower plastic substrate *W* . Together, these define a neural model that generates predictions about upcoming sensory inputs. Given an input stream {*x*_*t*_}, the model produces a one-step-ahead estimate 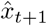 and incurs a prediction error *ϵ*_*t*_. The core NeuraGEM update rules (Eqs. 1–4 in Results) update both *Z* and *W* to reduce *ϵ*_*t*_, with *Z* being updated on a faster timescale. The gradients required for updating *Z* were computed using backpropagation through time (BPTT); a biologically motivated alternative at test time is described in section 3.7.

This formulation does not depend on a specific network architecture. The plastic substrate *W* may parameterize a wide range of neural networks, and the fast substrate *Z* may correspond to a small population of neurons, the state of a recurrent network, or the output of a recurrent network. In the present work, we implemented *W* using a recurrent network to enable direct comparison with conventional RNN-based models.

#### 3.1.1 Modulatory influence of *Z* on *W*

We considered two forms by which *Z* modulates ongoing neural activity. In the *multiplicative* form, *Z* gates the pre-activation of the recurrent population. If the recurrent core has pre-activation *a*_*t*_, then the gated pre-activation *u*_*t*_ is:

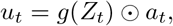

where the weights *g* mapping *Z* to the recurrent population were fixed and initialized from a Bernoulli distribution (probability 0.4). ⊙ denotes elementwise multiplication. Note that this pre-gating is applied prior to recurrent weights as opposed to post-gating.

In an *additive* form explored in neural analyses, *Z*_*t*_ was concatenated to the sensory input and projected through learnable weights. Both forms yielded similar qualitative behavior; multiplicative modulation was used unless noted otherwise.

### 3.2 Neural architecture, baselines, and optimization

#### 3.2.1 Neural network architecture

The plastic substrate *W* comprised three stages: (i) an input embedding multilayer perceptron (MLP), (ii) a recurrent neural population, and (iii) an output projection MLP. The recurrent population contained 32 units. We used an LSTM implementation as a stable and widely used recurrent network, particularly formulated for long input horizons [67], and to keep architectural comparisons consistent across models.

#### 3.2.2 Baseline recurrent models

We compared NeuraGEM to conventional recurrent networks trained on the same prediction objective. RNN^short^ and RNN^long^ differed only in the length of the input horizon used during training (see below) and did not include the fast neural substrate *Z*.

#### 3.2.3 Input horizon

At each time step, models generated a prediction of the subsequent observation. To examine online adaptation, networks updated *W, Z*, or both, before processing the most recent *h* observations again at the next time step. Short horizons typically include at most one latent switch, whereas long horizons may span multiple switches. NeuraGEM relied on short horizons, using fast updates of *Z* to adapt rapidly to changes in task structure.

#### 3.2.4 Training and Optimization

Models were trained using the Adam optimizer [29]. For NeuraGEM, we used distinct update rates (i.e., learning rates) for the fast substrate *Z* and the slower plastic substrate *W* (denoted *α*_*Z*_ and *α*_*W*_ in Results; see Eqs. 1–4). Adam’s momentum parameters were set to *β*_1_ = 0.5 and *β*_2_ = 0.7, which improved the responsiveness of fast *Z* updates while leaving overall training behavior unchanged.

All *Z* units were initialized to zero at the start of training. *Z* was regularized toward baseline using an L2 penalty implemented through the optimizer 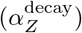.

Models were trained for 40 blocks in all experiments, and synaptic parameters were frozen for subsequent testing.

The multi-timescale experiment (fig. 5) was trained using stochastic gradient descent (SGD) rather than Adam to more directly isolate the effects of differences in the update and decay rates of distinct *Z* populations. Adaptive rescaling in Adam can reduce the contrast between different update and decay rates.

#### 3.2.5 Summary of task-specific hyperparameters

Key hyperparameters by task are listed in table 1.

**Table 1.** Hyperparameters by task. CST: contextual switching task. Update rates correspond to the coefficients in the NeuraGEM update rules reported in Results (Eqs. 1–4).

| Task | Model | horizon $h$ | $\alpha_Z$ | $\alpha_{\text{decay}}^Z$ | Optimizer |
| --- | --- | --- | --- | --- | --- |
| CST | NeuraGEM | 5 | 0.5 | $10^{-4}$ | Adam |
| CST | RNN<sup>short</sup> | 5 | – | – | Adam |
| CST | RNN<sup>long</sup> | 50 | – | – | Adam |
| EM demo | NeuraGEM | 5 | 0.5 | $10^{-4}$ | Adam |
| Seq. learn | NeuraGEM | 6 | 0.2 | $3 \times 10^{-5}$ | Adam |
| Seq. learn | RNN<sup>short</sup> | 6 | – | – | Adam |
| Seq. learn | RNN<sup>long</sup> | 200 | – | – | Adam |
| Multi-timescale | $Z_{\text{fast}}$ | 5 | 1.0 | 0.3 | SGD |
| Multi-timescale | $Z_{\text{slow}}$ | 5 | 0.5 | 0.15 | SGD |

### 3.3 Contextual Switching Task

Observations *x*_*t*_ were sampled from a Gaussian distribution whose mean alternated between 0.2 and 0.8 in blocks (*σ* = 0.3). Block sizes were sampled from a Geometric distribution with a mean of 25 and truncated to 15–50 time steps. Each model predicted the next observation at every time step (sequence-to-sequence training). Training spanned 40 blocks, after which weights were frozen and the models were tested under three manipulations:

- **Novel block sizes:** blocks shorter or longer than the training range.
- **Novel latent means:** Gaussian means varied from –0.2 to 1.2 (step 0.1).
- **Novel noise levels:** standard deviation *σ* increased beyond the training value.

Each condition was repeated across *n* = 20 random seeds. Whole block MSE is an average of all prediction errors in a block. Asymptotic (“block-end”) MSE was computed from the last 10 steps of a block.

#### 3.3.1 Behavioral learning rate

The behavioral learning rate quantifies how strongly the model updates its output in response to the previous prediction error:

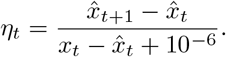

#### 3.3.2 Bayesian observer

An ideal Bayesian observer was implemented to benchmark inference. Observations were generated as

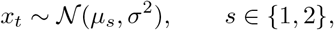

with latent state *s* switching according to a hazard rate *ρ*. The posterior was updated recursively:

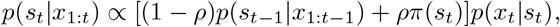

and normalized. The predictive mean was 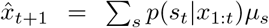. Distance to Bayesian predictions was computed as the mean absolute error across time.

### 3.4 Multi-timescales task

We extended the contextual switching task to a five dimensional input, also sampled from independent Gaussian distributions at each time step. Five latent variables determined the mean value of the Gaussian distribution at the corresponding input dimension. The latent variables switched at distinct rates, with block sizes sampled from Geometric distributions with means of 20, 60, 80, 120, and 160. In determining the correlation with Z units in the model, slower signals inherently had higher correlation. Therefore, we computed a null correlation value for each latent variable by computing the mean correlation of each latent variable with 200 circularly shifted versions of its activity.

### 3.5 Sequence learning task

We implemented the sequence learning paradigm from [39]. Six-step “stories” were generated by one of two latent Markov chains (contexts). Each possible story state was represented as a one-hot encoded vector.

#### Training curricula

Three curricula were used: (1) blocked (context constant for 40 stories), (2) interleaved (context alternates every story while keeping the same number of stories), and (3) mixed (interleaved followed by blocked).

#### Quantification

To quantify performance, following the procedure used in the experiment in humans [39], we transitioned models to a random curriculum phase in which the context was sampled randomly for each story over 80 stories. Importantly, this manipulation changes the temporal statistics of context. We hypothesize that humans can take advantage of cues signaling trial boundaries (e.g., an inter-trial interval) to decay internal states between trials, which will help in this random curriculum. However, NeuraGEM receives no such signals, and residual *Z* values from the previous sequence carry over at the start of each new sequence. To enable NeuraGEM to adapt to random context changes, we increased the update and decay rates of *Z* selectively during this random phase: 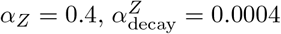, and *Z* was updated five times per time step.

Accuracy was measured as correct predictions on the third transition (choice between states 5 vs. 6), which is diagnostic of learning the correct computation [39]. The relationship between *Z* activity and the context-identifying transition (*T*_cid_) or random transition (*T*_rnd_) was quantified using the coefficient from a linear regression model.

### 3.6 Neural analyses

To relate model representations and dynamics to neural signatures reported in the literature, we performed three complementary analyses on hidden-unit activity.

#### 3.6.1 Cross-condition generalization performance (CCGP)

To assess whether context was represented in a stimulus-invariant manner, we computed cross-condition generalization performance (CCGP) using a linear support vector machine (SVM) [9]. For each model and random seed, stimulus conditions were defined *within each latent context* by splitting observations according to whether the instantaneous input was above or below that context’s mean value. This procedure ensured that the stimulus condition and latent context were not confounded.

The classifier was trained to decode context using hidden-unit activity from one within-context stimulus condition (e.g., below-mean inputs) and was tested on the complementary condition (above-mean inputs), and vice versa. The two test accuracies were averaged to obtain a single CCGP value per seed. Chance performance is 0.5. For NeuraGEM, only hidden-unit activity was used in the analysis; the activity of *Z* was excluded.

#### 3.6.2 Cosine similarity across context epochs

To quantify representational separation between contexts, hidden-unit activity was averaged across time steps within each context block. Cosine similarity was computed between successive blocks, yielding a measure of representational overlap (lower values indicate greater separation). For each seed and model, similarity values were averaged over early training blocks.

#### 3.6.3 Structure in Z to W projections

Hidden units were partitioned by context preference by comparing their mean activity across contexts. Units with higher average activity in Context A or Context B were assigned to the corresponding group. For the additive NeuraGEM variant, in which projections from *Z* to the recurrent population were learnable, we tested for asymmetries in these projections. For each seed and context block, the component of *Z* with the highest activity was identified, and its outgoing weights were averaged separately over units preferring each context.

#### 3.6.4 Attractor dynamics

To characterize autonomous dynamics, each trained network was treated as a discrete dynamical system and evolved without external input. Hidden-unit activity was projected onto the first two principal components for visualization. PCA was fitted to the testing phase data, per model. Fixed points were identified as states where the norm of the state update fell below a small threshold for three consecutive time steps.

### 3.7 A learned feedback projection replaces backpropagation

We introduced a feedback network trained to *predict* the appropriate update to *Z* directly from the instantaneous output residual and current *Z* activity (fig. 8A). The forward model computes

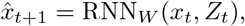

The feedback network RNN_*f*_ (8 units) maps the error incurred and the current *Z* activity to a gradient update:

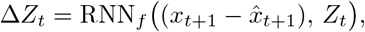

and *Z* is then updated:

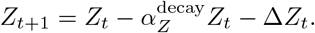

During training, RNN_*f*_ was supervised to approximate the gradient direction that would be obtained by backpropagating the prediction error through the forward network:

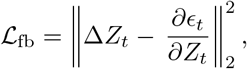

Where 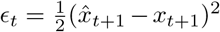. Gradients *∂ϵ*_*t*_*/∂Z*_*t*_ were computed using backpropagation through time during training only and served solely as teaching signals for RNN_*f*_ .

#### Reporting and multiple comparisons

All *p*-values are reported uncorrected. Given the small number of planned comparisons in each analysis, no multiple-comparisons correction was applied.

#### Significance code

∗ *p <* 0.05, ∗∗ *p <* 10^−2^, ∗ ∗ ∗ *p <* 10^−3^, ∗ ∗ ∗∗ *p <* 10^−4^; n.s. = not significant.

## Acknowledgements

We thank Matthew Nassar and Robert Guangyu Yang for their support and insightful discussions. This research was funded by the NIH grant R01-MH132172.

## 4 Appendix

### 4.1 Derivation of NeuraGEM from EM algorithm

This section provides a derivation of the gradient-based updates described in the main text and shows how NeuraGEM can be understood as a generalized EM algorithm under a delta variational approximation.

We begin by setting up the complete-data model. For a sequence *X* = (*x*_1_, …, *x*_*T*_) with latents *Z* = (*Z*_1_, …, *Z*_*T* −1_), the complete-data model factorizes as

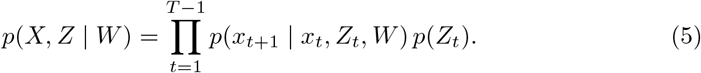

We assume a Gaussian observation model

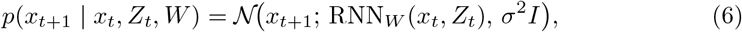

and a Gaussian prior on *Z*_*t*_

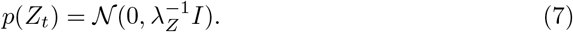

Up to additive constants, the complete-data log joint is

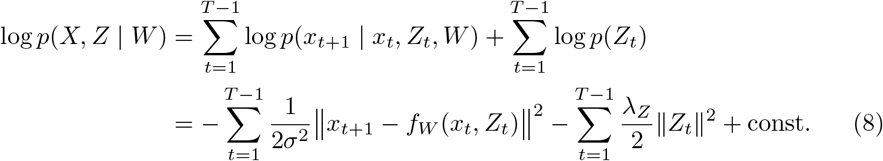

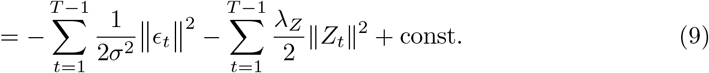

The goal is to maximize the marginal likelihood log *p*(*X*|*W*). However, this is usually intractable. We thus consider the following derivation of the evidence lower bound (ELBO). Let the probability distribution of **Z** be *q*. Specifically, instead of maintaining the whole distribution, NeuraGEM maintains a point estimate of *Z* by choosing a delta variational family

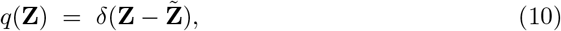

which reduces the ELBO to the complete-data log posterior.

We can write

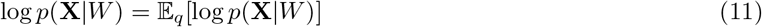

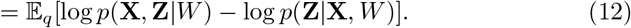

By adding and subtracting an entropy of *q*, we have

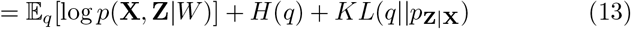

where KL divergence *KL*(*q*||*p*) is defined as 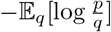 and the entropy *H*(*q*) is defined as ™𝔼_*q*_[log *q*].

Define ELBO as

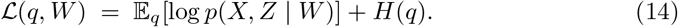

Because KL is non-negative, we have log *p*(**X**|*W*) ≥ ℒ (*q, W*). The equality only holds when KL is 0 and hence only when *q* = *p*_*Z*|*X*_ . In that case, we have 𝔼_*q*_[log *p*(**X, Z**|*W*)]+ *H*(*q*) = log *p*(*X*|*W*). Then, optimal weights *W*^∗^ can be found by maximizing ℒ (*q, W*) with *q* = *p*_*Z*|*X*_ . Because entropy has no dependency of *W*, we have

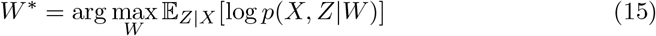

This recovers the standard EM coordinate-ascent formulation under exact posterior inference. Notice that this is equivalent to performing coordinate descent on ℒ(*q, W*) since maximizing ELBO with respect to *q* exactly occurs at *q* = *p*_*Z*|*X*_ .

NeuraGEM maintains a point estimate of the latent state, which corresponds to restricting the variational family to delta distributions 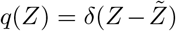. Since *H*(*δ*) = 0, the ELBO reduces to the complete-data log joint,

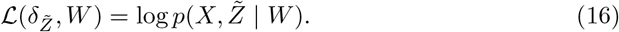

Maximizing 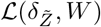 by coordinate ascent yields the following EM variants:

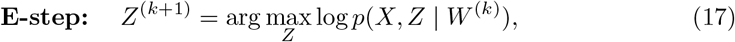

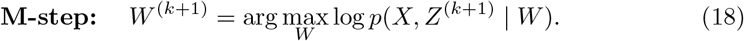

Exact maximization in Equation 17-18 is intractable for neural networks, so NeuraGEM performs generalized EM by taking a gradient step in each block.

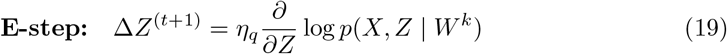

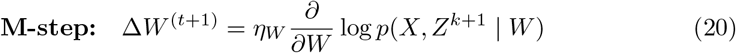

with *η*_*q*_ ≫ *η*_*W*_ so that *Z* effectively stays close to the optimal latent assignment whenever *W* updates, similar to coordinate ascent.

Finally, substituting the log joint (9) into (19)–(20) recovers the NeuraGEM update rules (Equation 4 and 3) up to the choice of optimizer and constant rescaling of the step sizes.

